# Catechol-Containing Compounds are a Broad Class of Protein Aggregation Inhibitors: II. Rosmarinic Acid Potently Detoxifies Amylin Amyloid and Ameliorates Diabetic Pathology in HIP Rats

**DOI:** 10.1101/2020.12.13.873687

**Authors:** Ling Wu, Paul Velander, Anne M. Brown, Yao Wang, Dongmin Liu, David R. Bevan, Shijun Zhang, Bin Xu

## Abstract

Protein aggregation is associated with a large number of human protein misfolding diseases, yet FDA-approved drugs are currently not available. Amylin amyloid and plaque depositions in the pancreas are hallmark features of type 2 diabetes. Moreover, these amyloid deposits are implicated in the pathogenesis of diabetic complications such as neurodegeneration. We recently discovered that catechols and redox-related quinones/ anthraquinones represent a broad class of protein aggregation inhibitors. Further screening of a targeted library of natural compounds in complementary medicine that were enriched with catechol-containing compounds identified rosmarinic acid as a potent inhibitor of amylin aggregation (estimated inhibitory concentration IC_50_ = 200-300 nM). Structure-function relationship analysis of rosmarinic acid showed the additive effects of two catechol-containing components of the RA molecule. We further showed that RA does not reverse fibrillation back to monomeric amylin, but lead to non-toxic, remodeled protein aggregates. Rosmarinic acid has significant *ex vivo* efficacy in reducing human amylin oligomer levels in HIP rat sera as well as in sera from diabetic patients. *In vivo* efficacy studies of rosmarinic acid treatment with the diabetic HIP rat model demonstrated significant reduction in amyloid islet deposition and strong mitigation of diabetic pathology. Our work provides new *in vitro* molecular mechanisms and *in vivo* efficacy insights for a model nutraceutical agent against type 2 diabetes and other aging-related protein misfolding diseases.

## Introduction

Protein misfolding diseases include a broad spectrum of neurodegenerative and related aging diseases such as Alzheimer’s disease (AD), Parkinson’s disease, prion disease, and type 2 diabetes (T2D) (Westermark et al, 2011; Eisenberg & Jucker, 2012; Knowles et al., 2014; Goedert, 2015; Chiti & Dobson, 2017). The pathological hallmarks of this class of diseases are amyloid fibrils: structurally conserved intracellular and extracellular insoluble proteinaceous deposits (Eisenberg & Sawaya, 2017). Amyloid formation proceeds through a nucleation-dependent aggregation process. Monomeric and oligomeric aggregates form amyloid “seeds” that initiate a cascade that results in equilibrium between mature amyloid fibrils and their small precursor aggregates (Holmes et al., 2014; Knowles et al., 2014). Over the last two decades, increasing evidence has indicated that the primary pathological amyloid species are these small non-fibrillar precursor aggregates, ranging from unstructured oligomers to β-sheet rich aggregates or protofibrils (Bieschke et al, 2011; Kayed & Lassagna-Reeves, 2013; Knowles et al, 2014; Chiti & Dobson, 2017).

Amylin, also referred to as islet amyloid polypeptide or IAPP, is a 37-amino acid peptide hormone co-secreted with insulin by pancreatic β-cells (Westermark et al, 2011; Cao et al, 2013). Physiologically, amylin plays important roles in glycemic regulation (Roberts et al, 1989; Westermark et al, 2011). Amylin is one of the most cytotoxic amyloidogenic proteins known, even more toxic than the well-studied amyloid β-peptide, Aβ42 (Abedini et al, 2013; Figs. S1A and S1B). A state of hyperamylinemia is associated with the compensatory increased insulin secretion that occurs during the development of insulin resistance and T2D due to the co-secretion of amylin and insulin hormones. Toxic amylin oligomer/amyloid formation and plaque deposition in the pancreas are hallmark features of T2D (Westermark et al, 1972; Verchere et al, 1996; Hoppener et al, 2000; Westermark et al. 2011). Experimental evidence shows that hyperamylinemia also induces toxicity in other organs, including the heart, kidneys, and the brain (Gong et al, 2007; Jackson et al, 2013; Srodulski et al, 2014; Verma et al, 2016). The accumulation of amylin is associated with vascular and tissue damage to the heart and brain (Despa et al, 2012; Jackson et al, 2013). While the rodent amylin sequence is not amyloidogenic (Westermark et al, 2011; Velander et al, 2020), a “humanized” transgenic diabetic rat model overexpressing human amylin (HIP rat) has provided strong causal evidence that amylin oligomers/amyloid contribute to heart dysfunction and AD-mimicking neurological deficits in addition to diabetes (Matveyenko & Butler, 2006a; Despa et al, 2012; Despa et al, 2014; Srodulski et al, 2014; Ly et al, 2017). Amylin oligomers were detected in the serum of HIP rats and the accumulation of amylin deposits in the hearts and the brains of these animals was also observed (Srodulski et al, 2014; Ly et al, 2017).

Currently, there are no cures for any protein amyloid diseases. Progress toward managing protein misfolding diseases in general has been hampered by the failure to develop effective disease-modifying drugs. This is partly due to our limited understanding of amyloidogenic proteins as well as their interactions with small molecules/inhibitors. Identification of effective amyloid inhibitors is challenging because of the intrinsic structural disorder of the protein targets of amyloid assembly; however, there are recent significant advances in the structural biology research of amyloid proteins, such as cryogenic electron microscopy (cryo-EM) characterization of their key fragments, which may enable structure-based inhibitor design in the future (Tycko, 2015; Fitzpatrick et al, 2017; Krotee et al, 2017; Eisenberg & Sawaya, 2017; Seidler et al, 2018; Falcon et al, 2018).

There are multiple therapeutic strategies for identifying disease-modifying agents for neutralizing toxic protein amyloids (Jiang et al, 2013; Eisele et al, 2015; Habchi et al, 2016). One of the current strategies aimed at identifying lead compounds focuses on inhibiting amyloid aggregation by (i) inhibiting toxic amyloid formation or stabilizing its native form from further aggregation and (ii) remodeling or degrading toxic amyloid oligomers and/or insoluble fibrils. For natural product-based amyloid inhibitor identification, one source of information from epidemiological studies suggests preventative effects against neurodegenerative dementia or diabetes may be associated with diets containing a high intake of flavonoids and polyphenolic compounds (Yamada et al, 2015; Velander et al, 2017). The importance of diet has been well recognized as dietary risks are ranked as one of the highest risk factors in aging diseases (Murray et al, 2013). Past epidemiological work as well as alternative medicine studies led to testable hypotheses and experimental efforts that have successfully identified numerous natural compound amyloid inhibitors (Ono et al, 2004; Rigacci et al, 2010; Ono et al, 2012; Ardah et al, 2015; Velander et al, 2016; Wu et al, 2017; Velander et al, 2020).

In order to gain insight into general classes of chemical structures of amyloid inhibitors, we recently screened an NIH Clinical Collection (NIHCC) drug library of 700 small molecule compounds with diverse chemotypes against three prototype amyloidogenic proteins amylin, Aβ, and tau. We identified catechols and redox-related quinones/anthraquinones as a broad class of protein aggregation inhibitors (Velander et al, 2020). To identify new, potent small molecule inhibitors of amylin amyloids, we further screened a targeted library of natural products used in alternative medicine enriched with catechol-containing compounds. This effort identified rosmarinic acid (RA), a catechol-containing natural product, as a potent inhibitor. This dual catechol-containing molecule demonstrated an additive effect, attributing the activities of the molecule to two active components, caffeic acid (CA) and salvianic acid A (SAA). Our inhibitor-induced amyloid remodeling data provided further evidence suggesting that amyloid remodeling is rapid and uses a pathway different from insoluble amyloid dissolution or fibril to monomer disaggregation. RA showed *ex vivo* efficacy in reducing human amylin oligomers in the sera from transgenic diabetic HIP rats and from that of diabetic patients. *In vivo* animal studies with the HIP rats demonstrated that RA potently reduced amyloid islet deposition and mitigated HIP rat diabetic pathology. Our work provides new *in vitro* molecular mechanisms and *in vivo* efficacy insights for a model nutraceutical agent against T2D and other aging-related protein misfolding diseases.

## Results

### Identification of Rosmarinic Acid

We previously screened a NIHCC drug-repurposing library of drugs and investigational compounds of diverse chemical structures diverse and discovered that catechol-containing compounds are a broad class of amyloid inhibitors (Velander et al, 2020). Based on this finding, we further rapidly screened a small library of natural compounds (~100 compounds) that are enriched in catechol-containing compounds used in Complementary Medicine for their anti-diabetic, anti-inflammatory, or neuroprotective effects. Epigallocatechin gallate (EGCG) was used as a positive control as it is extensively reported in literature for its strong anti-amyloid effects (Figs. 1A & 1B; Ehrnhoefer et al, 2008; Meng et al, 2010; Pithadia et al, 2016). Several other compounds including morin, baicalein, salvianiolic acid B, resveratrol, caffeic acid, and oleuropein were confirmed in our screen for their inhibitory effects against amylin amyloid formation (Fig. 1A; Cheng et al, 2011; Noor et al, 2012; Cheng et al, 2013; Tu et al, 2015; Velander et al. 2016; Wu et al, 2017). RA was discovered to be one of the highest-ranking inhibitors in blocking amylin amyloid formation (Fig. 1A). We further validated RA in multiple secondary assays. Using transmission electron microscopy (TEM) analysis, we found that RA significantly reduced amyloid fibril formation (Fig. 1B; estimated to be 50%), while at the same time enhancing the formation of non-toxic aggregates or amorphous aggregates at a 3:1 drug:amylin molar ratio (Fig. 1B). This potency is similar to the control compound EGCG. Based on thioflavin T (ThT) fluorescence assay, we further determined that the inhibitory concentration IC_50_ for RA is at 200-300 nM, several folds stronger than EGCG (estimated to be 1 μM) (Fig. 2A). Consistently with these data, RA also neutralized amylin amyloid induced cytotoxicity in pancreatic INS-1 cells and neuronal Neuro2A cells (Fig. 2B). RA further significantly delayed amylin amyloid formation. This was evidenced by RA causing a dose-dependent kinetic delay in the observed t1/2 values (Fig. 2C).

**Figure 1.**
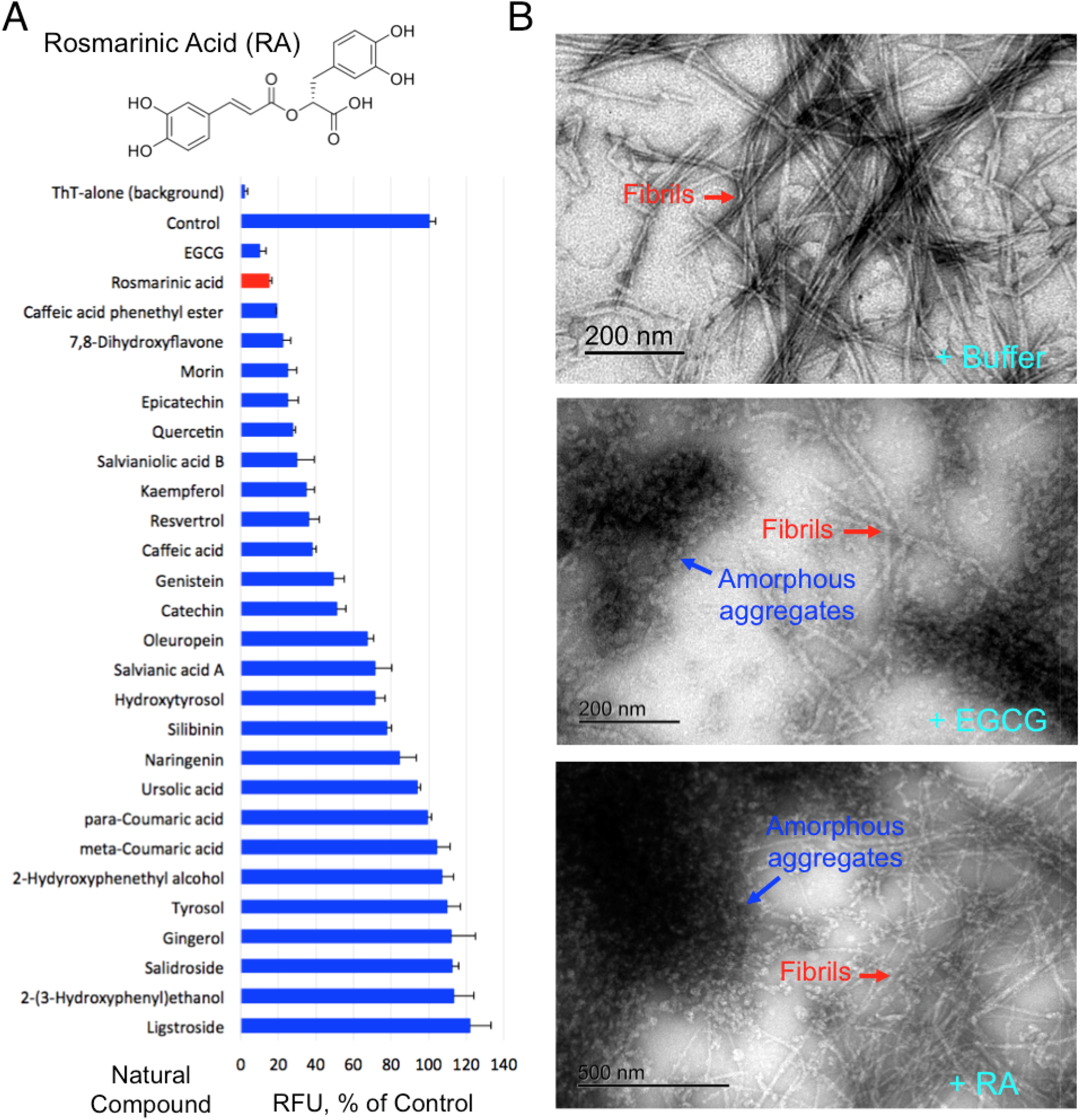
Identification of RA as a potent amylin amyloid inhibitor. (A) Identification of rosmarinic acid as strong amylin amyloid inhibitor from a natural product library enriched with catechol-containing compounds, a class of broad amyloid inhibitors we recently discovered (Velander et al, 2020), using a ThT fluorescence based screening. EGCG was used as a positive control. Amylin concentration was 5 μM and molar ratios of compound to amylin were 2:1. Chemical structure of RA is shown. (B) TEM images of human amylin amyloid with and without inhibitor treatment (1:20 amylin:drug molar ratio). Mature fibrils and amorphous aggregates are indicated by the red and blue arrows respectively. Buffer treatment sample served as control (100% mature fibril). RA treatment resulted in estimated 50% of mature fibrils and 50% of amorphous aggregates whereas treatment of EGCG, a known strong inhibitor, resulted in similar partial fibrils and partial amorphous aggregates.

**Figure 2.**
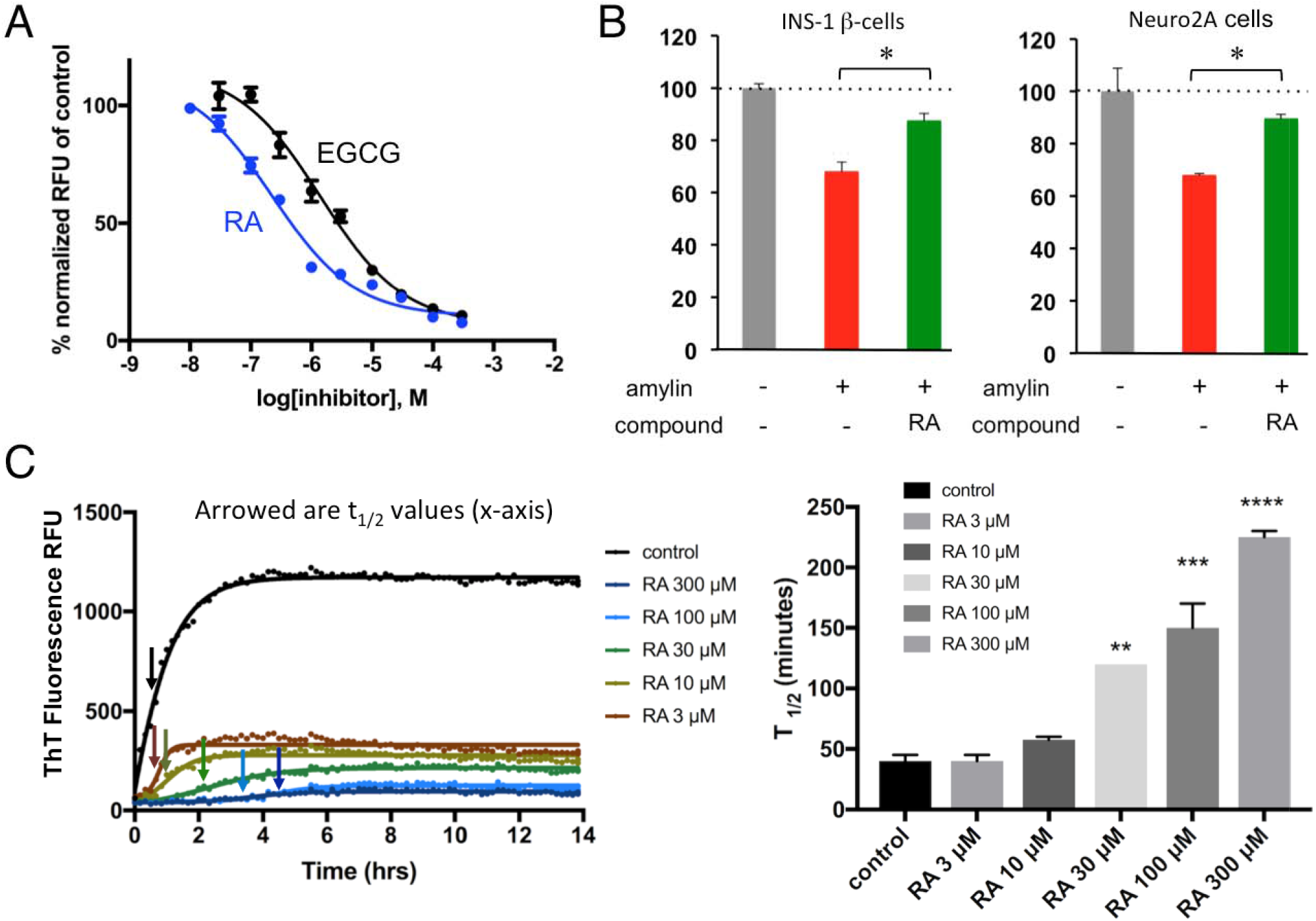
Characterizations of RA as a strong amylin amyloid inhibitor. (A) Amylin-ThT fluorescence based inhibition assay. RA is estimated to have IC_50_ between 200-300 nM. IC_50_ curve for control EGCG is also shown (1 μM) as a comparison. (B) Neutralization of amylin amyloid-induced cytotoxicity by RA in pancreatic INS-1 cells and neuronal Neuro2A cells. Amylin concentration was 3.75 μM and the ratio of inhibitor:amylin was 5:1. RA significantly detoxified amylin amyloid-induced cytotoxicity in both cell lines, as indicated by asterisks (p<0.05). (C) RA-induced dose-dependent kinetic delays in amyloid formation. Left panel shows the measurements of t1/2 of amylin amyloid formation by amylin-ThT fluorescence based assays, as indicated by the arrows. Concentration of amylin was 10 μM. Time points for t_1/2_ are shown as arrowed. At least three repeats have been performed. Right panel shows corresponding quantification and display in bar graph format of t_1/2_ values with each treatment. With increased concentration of RA treatment, t_1/2_ becomes significantly delayed (** p<0.05; *** p<0.01; **** p<0.001).

### Additive Effects of Rosmarinic Acid

Because RA consists of two catechol-containing components, caffeic acid (CA) and salvianic acid A (SAA) with a link of ester bond (Fig. 3A), we hypothesized that each component, CA and SAA, would also inhibit amyloid formation and that their added effects would be similar to that of RA. To test this hypothesis, we first performed ThT fluorescence assays, in which CA and SAA both showed significant amyloid inhibition activities, but neither as potently as RA. A mixture of CA and SAA at a 1:1 molar ratio showed inhibitory activity similar to that of RA (Fig. 3A). We further quantitatively determined the IC_50_ values for RA, CA and SAA as 200-300 nM, 7.2 μM, and 90 μM, respectively (Fig. 3B). We also investigated the remodeling capacity of RA, CA, SAA, and a mixture of CA and SAA after the ThT fluorescent signal of amylin amyloid reached plateau. Rapid drops in ThT signals were observed when preformed amylin amyloids were spiked with RA, CA, SAA, and CA+SAA (a mixture of CA and SAA in equimolar ratio) (Fig. 3C).

**Figure 3.**
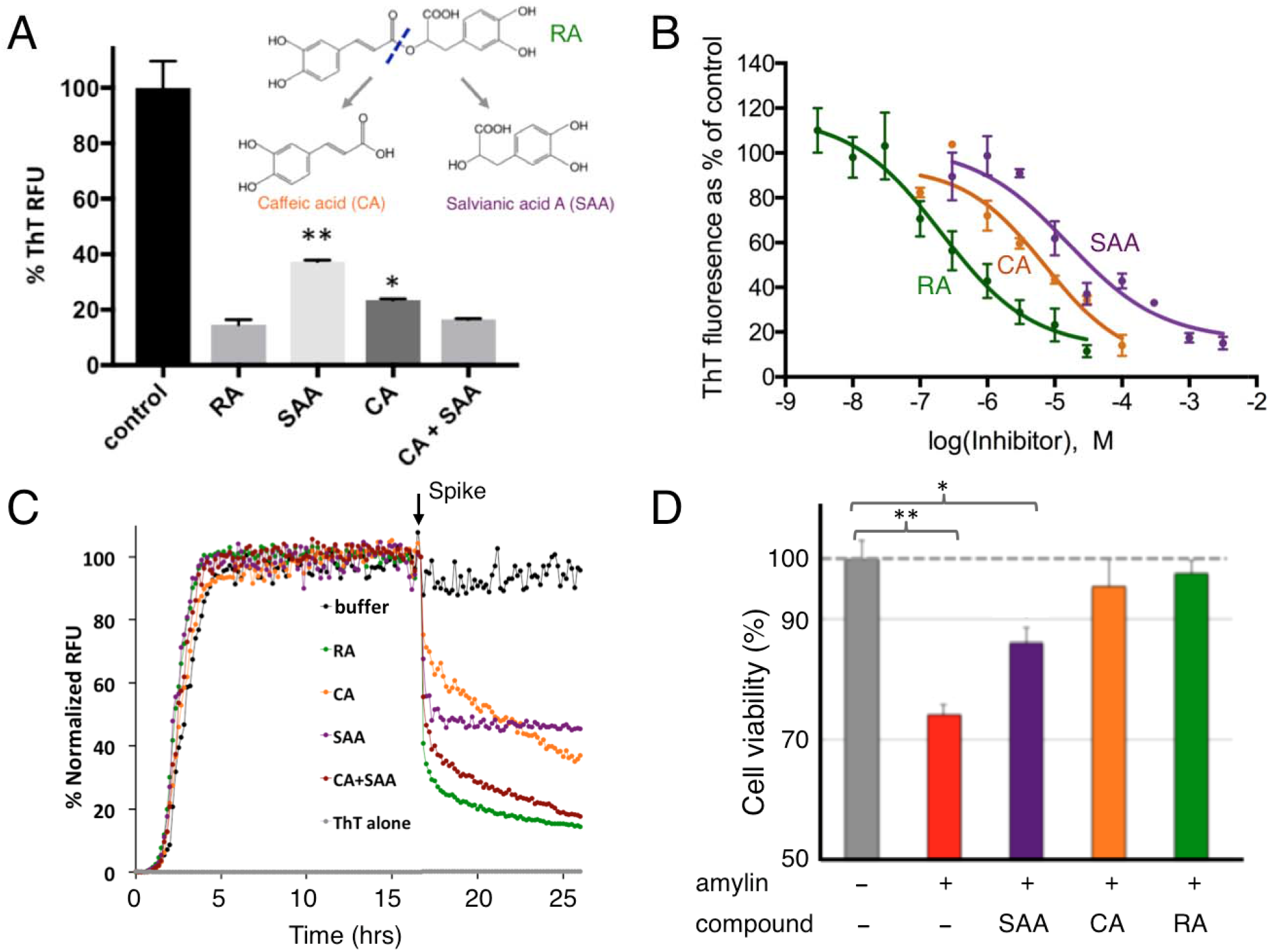
Additive effects of RA components in amylin amyloid inhibition. (A) ThT fluorescence based amyloid inhibition assay by RA and its hydrolytic components CA and SAA. Inhibition from 1:1 molar ratio of mixture of CA and SAA is similar to that of RA, but significantly stronger than those from CA or SAA alone. (B) IC_50_ measurements by ThT fluorescence based inhibition assays. IC_50_ is estimated to be 200-300 nM for RA, 7.2 μM for CA, and 90 μM for SAA. (C) ThT fluorescence-based amyloid remodeling assay shows amyloid remodeling after spiking of buffer control, RA, CA, SAA, and CA+SAA (1:1 molar ratio) after aggregation reaches plateau (t=17 hours). (D) Neutralization of amyloid-induced toxicity by RA, CA, and SAA in Neuro2A cells. Amylin concentration was 2.5 μM and the ratio of the inhibitor:amylin is 5:1. Comparing to the control, RA, and to lesser degrees, CA and SAA, significantly neutralize amyloid-induced toxicity. *, p<0.05, **, p<0.01.

The relative amyloid remodeling potencies depicted by these data are consistent with the anti-amyloid additive effects demonstrated by RA, CA, and SAA (Fig. 3A). This additive effect was also manifested in amylin amyloid-induced cytotoxicity rescue assays. At the inhibitor:amylin ratio of 3:1 (amylin concentration of 15 μM), RA fully neutralized amylin-induced neurotoxicity while CA and SAA only partially rescued the cell viability (Fig. 3D).

In parallel with biochemical and cell biology evaluation, we performed molecular dynamics (MD) simulation studies to test the additive effect hypothesis. Simulations were performed using fragments of amylin consisting of residues 20-29; this fragment has been shown to be the core sequence that drives fibril formation (Westermark et al, 1990; Cao et al, 2013). We first confirmed that these fragments form distinct β-sheet structures during MD simulation that mimic experimental nuclear magnetic resonance studies (PDB access code: 2KIB). We performed simulations using three human amylin (20-29) fragments in the absence of any inhibitor and in the presence of either RA, CA, or SAA (5 molecules inhibitor/system). Significantly less β-strand structure was observed in the simulations in the presence of RA (28 ± 11%) in comparison with no drug control (58 ± 11%). The amount of β-strand structure was also reduced in the presence of CA (37 ± 17%) and SAA (45 ± 20%), though not to the extent of RA (Table S1). Therefore, the MD studies showed collaborative results in terms of effectiveness in preventing aggregate formation as the experimental studies. Furthermore, analysis of inter-peptide interactions revealed a reduction between peptide fragments in the presence of RA, CA, and SAA as evaluated from average hydrogen bond presence (Table S2). Consistently, the reduction of interactions was more pronounced in the case of RA than with both CA and SAA, as represented by the non-diagonal peaks in the peptide fragment interaction “heat maps” (Fig. 4). Reduction of amylin fragment b-strand stacking is also evidenced by increased solvent accessible surface areas of amylin fragment in the presence of RA, CA, or SAA (Table S3). Collectively, these molecular simulation studies provided atomic-level mechanistic insights into the interactions between RA, CA, SAA and the amyloidogenic fragment (residues 20-29) of amylin. These findings also were consistent with our experimental results that demonstrated the additive effect of RA described above. Additionally, these MD studies provided us with a model system to study amylin aggregation and the effect of small molecules on these pathways, as well as a correlate at the atomistic level compared to assays performed in this study.

**Figure 4.**
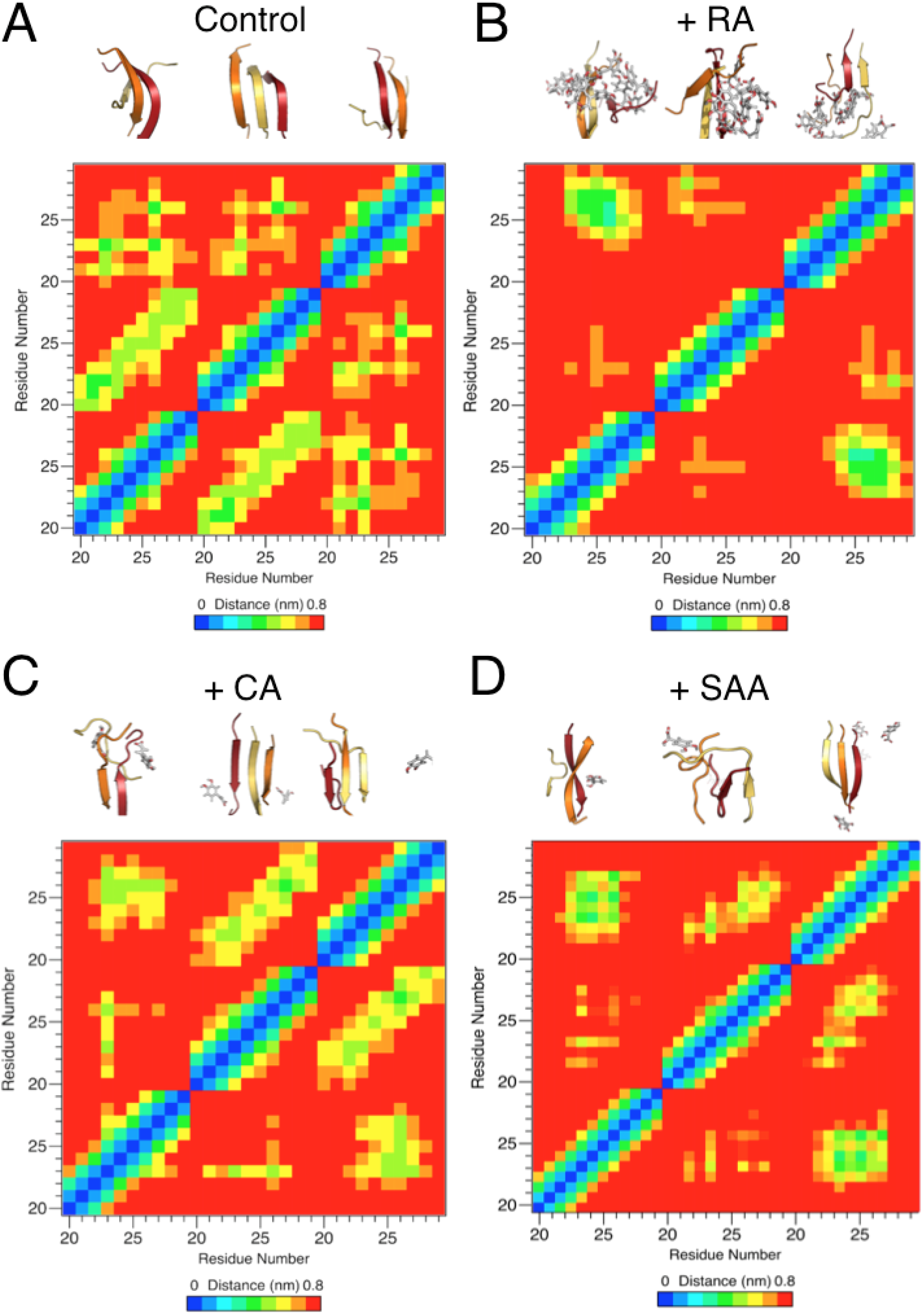
Additive effects of rosmarinic acid components in amylin amyloid inhibition with molecular simulation studies. Dominant morphologies of amylin_20-29_ trimer formation with and without the presence of RA, CA, and SAA are shown in the upper portions of each panel. Individual amylin_20-29_ peptide fragments are shown in cartoon and colored red (peptide 1), orange (peptide 2) and yellow (peptide 3). RA, CA, and SAA are shown as grey sticks, colored by element. For the panels of CA and SAA, only CA and SAA molecules within 1.0 nm of the amylin_20-29_ trimer are shown for clarity. Significantly less β-structure is observed in the presence of RA and reduction in β-structure is observed in the case of CA and SAA. Residue-residue interaction plots for corresponding drug treatment are shown as “heat” maps in the lower portions of each panel. X- and y-axis labels are in peptide fragment length (residues 20-29), with each repeating number (e.g., 20) being a separate peptide. Significantly less inter-molecular interactions between amylin fragments is observed in the presence of RA in comparison with the control. Reduction in interactions is observed in the cases of CA and SAA treatment.

### Inhibitor-Induced Amyloid Remodeling

RA demonstrated significant amyloid remodeling function similar to EGCG in orthogonal ThT fluorescence assays and gel-based remodeling assays (Fig. 5; Ehrnhoefer et al, 2008; Bieschke et al, 2010; Palhano et al, 2013). Importantly, RA neutralized amylin amyloid-induced cytotoxicity (Fig. 3D) and itself had no significant effects on cell viability for a variety of cell lines (Fig. S2). RA remodeled amylin amyloid into non-toxic aggregates, presumably via off-pathway channels as exemplified by EGCG (Ehrnhoefer et al, 2008; Bieschke et al, 2010; Meng et al, 2010; Palhano et al, 2013). We found that mixing RA with amylin amyloid led to sodium dodecyl sulfate (SDS)-resistant, remodeled aggregates spanning broad range of molecular weight that are non-toxic, as has been observed with EGCG in the literature (Palhano et al, 2013; Figs. 5B-5D). Remodeled molecular species were experimentally captured in as quickly as 45 minutes (boxed in the red rectangle; Fig. 5B). Amylin amyloid remodeling may also be shown in ThT fluorescence assays. Spiking RA or EGCG rapidly remodeled the preformed amylin amyloid (Fig. 5A). The same general steep drop in ThT fluorescence was observed when RA and EGCG were spiked at the plateau phase of amyloid formation (t=17 hours) and after mature fibrils were formed (t=3 days). No significant fluorescent signals for amyloid formation were observed when these compounds were spiked in at t=0 or before the amyloid growth phase (boxed in blue rectangle; Fig. 5A).

**Figure 5.**
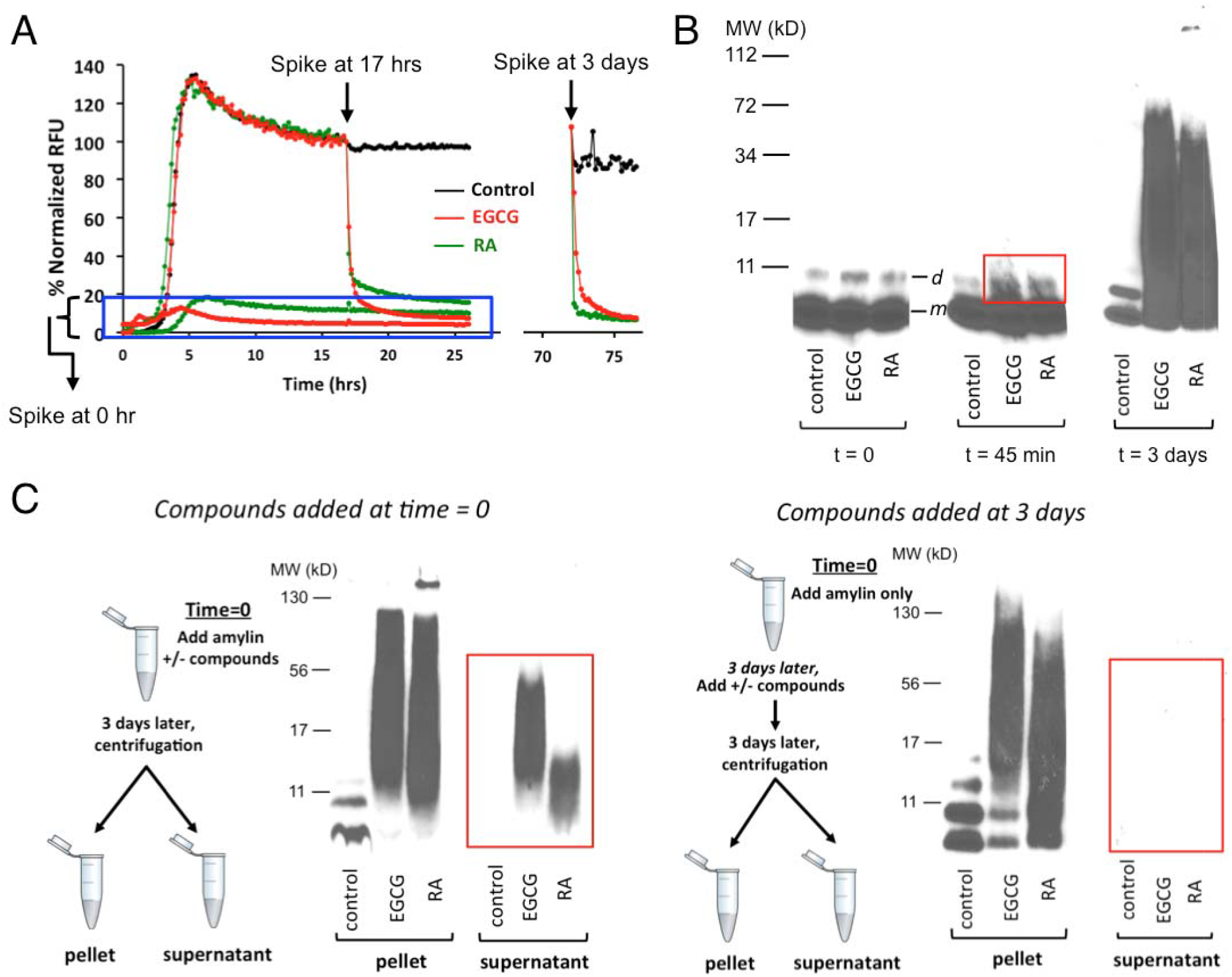
Amylin amyloid remodeling induced by RA as probed by ThT fluorescence based assays and orthogonal gel-based assays. (A) ThT fluorescence-based assay shows amylin amyloid remodeling after spiking of buffer control, RA, and EGCG before amylin amyloid is formed (t=0 hour), after amyloid has been formed (t=17 hours), or mature fibrils are formed (t=3 days). (B) Single fraction analysis of inhibitor-induced amyloid remodeling. Incubation of amylin amyloid with buffer control, EGCG, and RA leads to broad molecular weight range, SDS-resistant, remodeled protein aggregates. Samples were separated by SDS-PAGE and probed by an antibody specific for human amylin (T-4157). Urea-dissolved, broadly distributed molecular weight of amylin-inhibitor aggregates were observed while only monomer (indicated as “*m*”) and dimer bands (indicated as “*d*”) of amylin were seen in untreated samples. While with the quenching with denaturing conditions (6.5 M urea), both EGCG and RA induced broad molecular weight aggregates. SDS-resistant aggregates may be captured starting from 45 minutes of incubation (boxed in the red rectangle). (C) “Two-fractions” experiments demonstrated that EGCG and RA interact with monomeric amylin to form both soluble and insoluble aggregates of broad molecular weight range but such interactions with preformed fibrils lead to only insoluble aggregates. The experimental setting was the same as those described in panel 5B except that the pellet and the supernatant fractions (“two-fractions”) of each sample were separated by ultracentrifugation before the samples were separated by SDS-PAGE and probed by an antibody specific for human amylin (T-4157). Spiking of buffer control, EGCG, and RA was initiated either at 0 hour (left panel) or after mature fibrils were formed (t=3 days; right panel) as depicted in the schemes drawn on the left of each panel.

The mechanisms of remodeling by small molecule inhibitors and the physical basis of the related species changes and molecular interactions are not fully understood. The rapid drop in ThT fluorescence has been proposed to reflect disaggregation of insoluble mature amyloid fibrils in some cases (Meng et al, 2010). In other cases, this signal drop was proposed to reflect ThT binding site competition, altered stability, and seeding capacity of the remodeled fibril masses but not their solubility (Palhano et al, 2013).

Next, we used a “two-fraction” gel-based remodeling assay to assess if potential remodeling activities against preformed mature amyloid fibrils would result in their dissolution or disaggregation of fibrils to monomers. Herein, we spiked RA or EGCG with either freshly dissolved unaggregated amylin or with preformed mature amyloid fibrils, and after three days (sufficient time for mature fibril formation), analyzed the insoluble and soluble fractions from both conditions by Western blotting analysis using amylin-specific antibody T-4157 (Fig. 5C). As expected for amyloid inhibitors, all compounds maintained a significant amount of peptide mass within the soluble fraction (highlighted in the red rectangle; Fig. 5C left panel) when spiked into solutions initially containing freshly dissolved unaggregated amylin (i.e., a significant amount of insoluble amyloid formation was prevented). Moreover both RA and EGCG induced amyloid remodeling towards denaturant-resistant aggregates, whether compounds were spiked into solutions initially containing unaggregated amylin or preformed amylin fibrils (Fig. 5C). However, no compound was able to dissolve preformed amyloid fibrils nor did any compound disaggregate fibrils to amylin monomers, as reflected by the absence of any detectable amylin mass or monomeric amylin band in the soluble fraction of amylin fibril samples treated with the compounds (highlighted in the red rectangle; Fig. 5C right panel). Our data demonstrated that amylin amyloid remodeling by RA/EGCG is not simply the reverse pathway of amyloid formation; remodeling uses a pathway different from disaggregation of fibrils to monomers. Furthermore, both RA- and EGCG-induced amyloid remodeling showed differential solubility effects of remodeled off-pathway aggregates that is amyloid formation stage-dependent: EGCG and RA interact with monomeric amylin to form both soluble and insoluble aggregates of broad molecular weight range but such interactions with preformed fibrils lead to only insoluble aggregates (Fig. 5C), suggesting irreversibility once mature fibrils are formed. Conceivably, fibril formation may conceal certain RA interaction sites. To our knowledge, such differential solubility effects of inhibitor-remodeled aggregates have not been reported. This finding may have significant implication practically in selecting effective inhibitors to reduce pre-existing fibrils/plaques for therapeutics development for amyloid disease treatment.

### *Ex vivo* Efficacies of RA

Because rodent amylin is not amyloidogenic (Westermark et al, 2011; Velander et al, 2020), a “humanized” diabetic rat model (HIP rats), which overexpresses human amylin, has been established for mechanistic and translational applications (Butler et al, 2004; Srodulski et al, 2014). Past studies have confirmed that sera from HIP rats contain preformed amylin oligomers (Despa et al, 2012; Srodulski et al, 2014). Thus, as a step towards testing *in vivo* efficacy studies of RA, we obtained sera from HIP rats and then treated them with either RA or vehicle buffer control. These samples were then analyzed via Western blots using amylin-specific antibody T-4157. In contrast to vehicle-loaded controls, RA effectively reduced amylin oligomers (estimated to be 72 kD, 84 kD, and 96 kD) in a dose-dependent manner (Fig. 6A). Amylin oligomers of similar sizes from HIP rats have been reported in the literature (Srodulski et al, 2014).

To test whether RA has therapeutic potential for future usage in human, we investigated RA with sera from diabetic patients, as amylin oligomers had been detected in such clinical samples (Liu et al, 2016). We showed that RA effectively reduced amylin oligomers (estimated to be 52 kD) in sera from diabetic patients at a treatment concentration of 50 μM (Fig. 6B). As expected, reduction of amylin oligomer band intensity from diabetic patients was shown to be incubation time-dependent. The difference in molecular weights of the amylin oligomers detected between HIP rats and patient sera samples may be due to variation of amylin oligomer size in different hosts and/or different degrees of posttranslational modifications. Overall, RA was effective at the incubation time range of 3 hours to 24 hours and at concentration ranges of 10 - 50 μM in sera from both HIP rats and diabetic patients,

**Figure 6.**
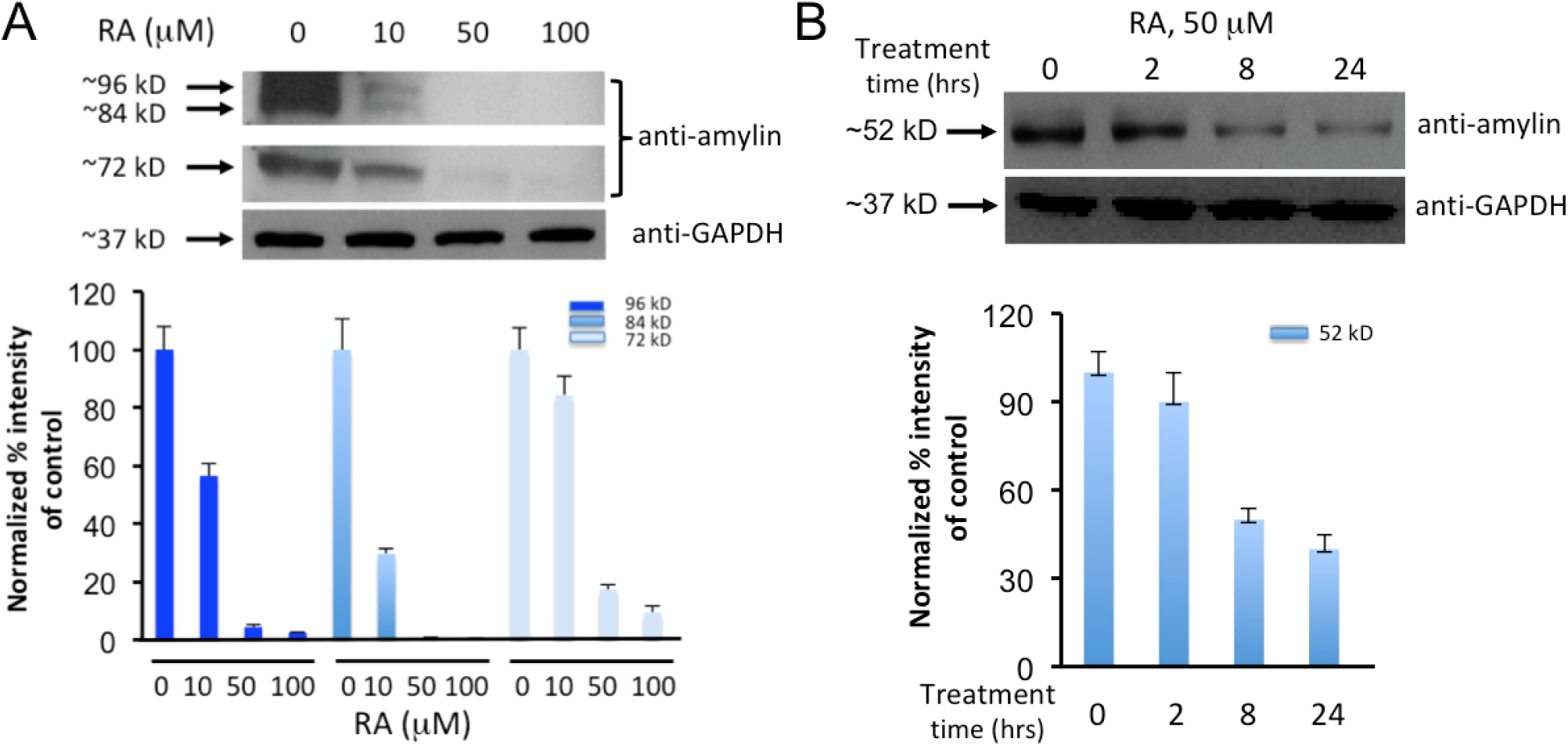
*Ex vivo* experiments demonstrate RA significantly reduces human amylin oligomers in the sera samples from HIP rats and diabetic patients. (A) Western blot analyses of HIP rat sera treated with increased concentrations of RA as indicated. Samples were incubation with RA at indicated concentrations for 3 hours at 37 °C. Equal amount of sera were loaded in each well as demonstrated by equal intensities of Western blot bands of control GAPDH. Lanes labeled as 0 μM are the no treatment controls. Anti-human amylin blots were probed with amylin specific antibody (T-4157). Estimated sizes of ~72 kD, ~84 kD, and ~96 kD of human amylin oligomers are indicated. Quantification of the Western blot results by densitometry are shown in the lower panel. (B) Western blot analyses of diabetic patient sera samples treated with 50 μM of RA as indicated over a time course of 0, 2, 8, and 24 hours. Equal amount of sera were loaded in each well as demonstrated by equal intensities of Western blot bands of control GAPDH. Anti-human amylin blots were probed with amylin specific antibody (T-4157). Estimated size of the human amylin oligomer (~52 kD) is indicated. Western blot results were quantified by densitometry, shown in the lower panel.

### *In vivo* Efficacy Studies of RA

HIP rats had been established as an excellent model to study amylin amyloid-induced islet pathology and to test novel approaches to the prevention and treatment of T2D and related complications (Matveyenko & Butler, 2006a; Despa et al, 2012; Srodulski et al, 2014). Hemizygous HIP rats spontaneously developed midlife diabetes (6-12 month of age) associated with islet amyloid (Butler et al, 2004). To test RA in reducing amylin amyloid oligomer formation in sera, pancreatic amyloid deposition, and in ameliorating islet diabetic pathology, we performed *in vivo* efficacy tests in hemizygous HIP rats with RA treatment via diet supplementation at a dose of 0.5% (w/w). The dose was chosen based on literature where several phenolic compounds (RA included) were tested in against Aβ amyloids in Tg2576 mice and showed *in vivo* efficacies (Hamaguchi et al, 2009). The dietary treatment was initiated when HIP rats were at prediabetic stage (at the age of 6 month; Butler et al, 2004) and the treatment lasted for four months. Due to significantly delayed phenotype in female rats, only male HIP rats were used (Ly et al, 2017). While EGCG is an excellent control for *in vitro* biochemical studies, it has not shown *in vivo* efficacy and failed in a recent clinical trial against α-synuclein aggregation (multiple system atrophy; Levin et al, 2019). Therefore, EGCG was not chosen to be included in our animal studies.

Our results demonstrated that RA showed highly effective anti-amyloid formation and anti-diabetic activities *in vivo* that were consistent with its *in vitro* activities described in this work: (1) As expected, diabetic HIP rats (10-12 month old) showed severe amylin amyloid deposition in the pancreatic islets as visualized by Congo Red staining (Fig. 7D, middle panel). The extensive amyloid islet deposition was reminiscent of a clinical thioflavin S-staining of the islets from a human with non-insulin dependent diabetes mellitus or T2D (Verchere et al, 1996). In contrast, wide type age-matched Sprague-Dawley (SD) rats showed no amyloid deposition in the islets (Fig. 7D, left panel). Amylin amyloid deposition in islets was significantly reduced in RA-treated, age-matched HIP rats (Fig. 7D, right panel). However, residual amyloid deposition was evident, suggesting that RA treatment regimen did not completely block amylin aggregation and deposition on the pancreas. (2) Immunohistochemistry analysis of islet tissues provided collaborative evidence that amylin deposition levels were significantly higher in the islets of untreated HIP rats in comparison with wild type SD rats and RA-treated HIP rats (Fig. 7E); (3) We also evaluated sera amylin oligomer levels by Western blots using amylin-specific antibody T-4157. In contrast to wild type SD rat controls that showed insignificant levels of amylin oligomers, untreated HIP rats showed strong intensities of amylin oligomers (estimated to be 32 kD and 52 kD respectively). With RA treatment, corresponding oligomer levels were either significantly reduced (52 kD bands) or fully abrogated (32 kD bands) (Fig. 7C). Amylin oligomers of similar sizes from HIP rats have been reported in the literature (Srodulski et al, 2014); (4) We further evaluated diabetic conditions of each cohort of rats. Wild type SD rats have an average non-fasting blood glucose level of 146.8 mg/dL. Untreated HIP rats have an average non-fasting blood level of 242.3 mg/dL that is evidently diabetic. Such spontaneous diabetes development in HIP rats and hyperglycemia are consistent with what have been reported (Butler et al, 2004; Srodulski et al, 2014). Noticeably, RA-treated HIP rats have an significantly reduced average non-fasting blood glucose level of 181.0 mg/dL (Fig. 7A). Consistently, serum insulin levels were significantly lower in untreated HIP rats than those in wild type SD rats and RA-treated HIP rats at the age of 10 months (Fig. 7B). Hypoinsulinemia state of HIP rats at the age of 10 months had been reported (Matveyenko & Butler, 2006b). We do not find any significant differences between RA-treated vs. non-treatment HIP groups on body weight and food intake (data not shown).

**Figure 7.**
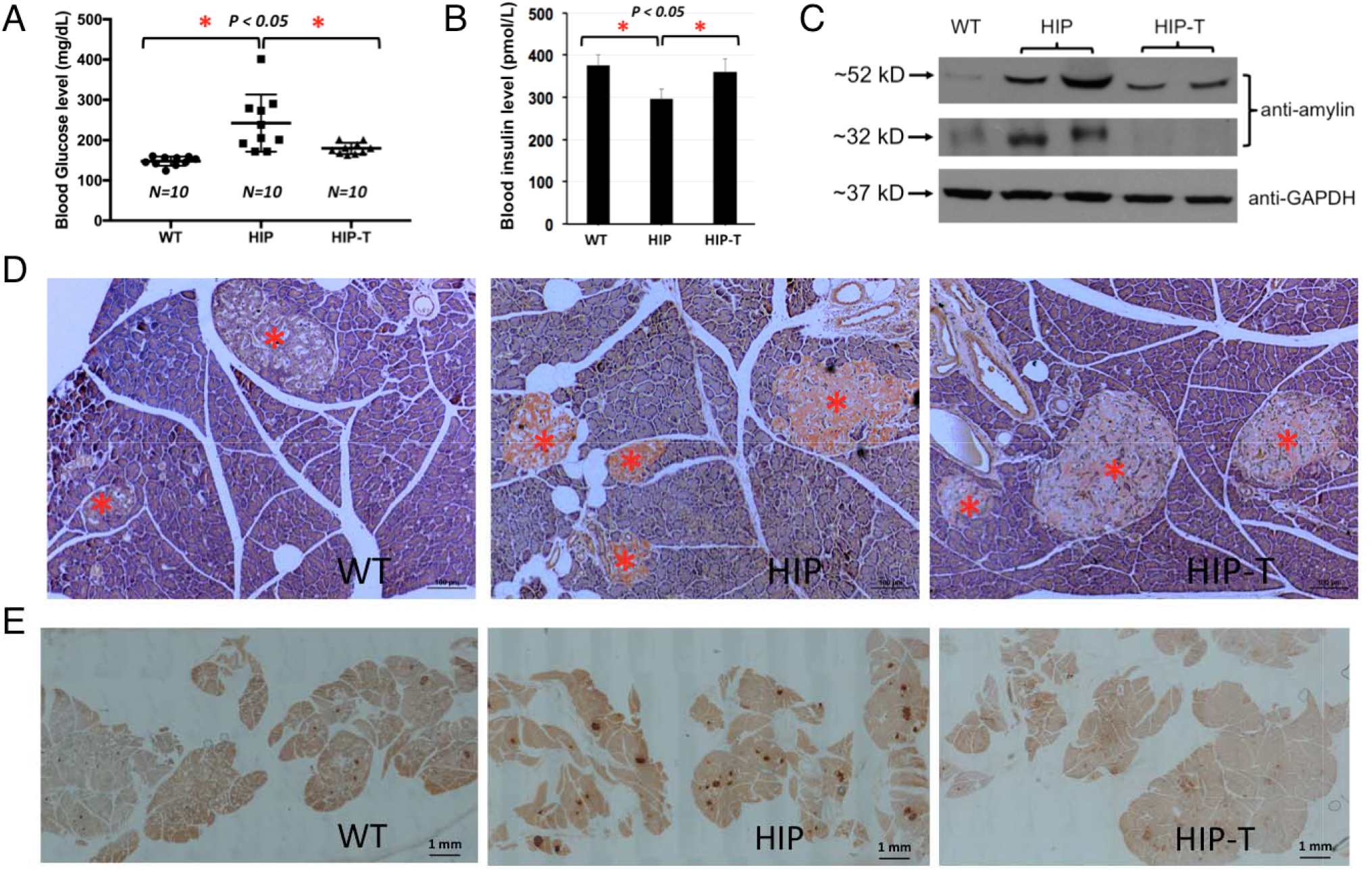
*In vivo* experiments demonstrate RA potently reduced amylin amyloid islet deposition and ameliorated diabetic pathology in HIP rats. Prediabetic HIP rats (6 month old) were treated (HIP-T) or untreated (HIP) with 0.5% w/w RA in regular chow diet for 4 months. Age-match wild type Sprague-Dawley rats with regular chow diet served as negative controls. At the end of the treatments, rats were euthanized and pancreatic tissues and sera were collected for immunohistochemistry and biochemical analysis. (A) Serum glucose levels were quantified for each group of rats. Significant reduction of serum glucose concentration was observed and indicated by asterisks (p<0.05) comparing the untreated HIP rats with those of RA-treated HIP rats and control SD rats. (B) Insulin levels were measured for each group of rats (n=4). Statistically lower levels of insulin was observed and indicated by asterisks (p<0.05) comparing the untreated HIP rats with those of RA-treated HIP rats and control SD rats. (C) Western blot analysis of serum amylin oligomer levels with amylin specific antibody (T-4157). GAPDH served as loading controls. Comparing with the untreated HIP rat sera, significant reduction of the 52 kD amylin oligomer bands and fully abrogated 32 kD oligomer bands were observed with RA-treated HIP sera. There were essentially no intensities of these oligomer bands in control SD rat sera. (D) Representative Congo red staining demonstrated strong amyloid staining in untreated HIP rat pancreatic tissue slices (orange-red color in the islets, middle panel). Significantly reduced Congo red staining in the islets of RA-treated HIP rats (right panel). No Congo red staining was observed in islets in control wild type SD rats (left panel). All pancreatic islets are marked by red asterisks. These qualitative results represent multiple samples from RA-treated or untreated HIP rats, or WT Sprague-Dawley controls (n=4) and at least 50 islets from pancreatic head, body and tail for each group were examined. Scale bar in each panel is 100 μm. (E) Representative amylin deposition in the pancreatic tissue slice using amylin antibody staining from different rat groups. Pancreatic tissue slices were immune-stained with amylin specific antibody (T-4157) followed by secondary antibody and NovaRED substrate (Vector Lab) treatment. Stained islets are shown in dark-red. Islets in the pancreatic slices from untreated HIP rats were intensely stained whereas those from RA-treated and control SD rats were only weakly stained. At least 10 tissue slices were observed from each group. Scale bar in each panel as indicated is 1 mm.

In summary, we demonstrated *in vivo* efficacy of RA in mitigating diabetic pathology and reducing amylin amyloid deposition in the pancreas of HIP rats. To our knowledge, this is the first report describing *in vivo* effectiveness of RA against amylin aggregation. We also discovered additive effects of its two active components, caffeic acid and salvianic acid A on amylin fibrillation by a combination of biochemical, cell-based and computational approaches. We showed that RA does not reverse fibrillation back to monomeric amylin, but lead to non-toxic, remodeled protein aggregates.

## Discussion

Several lines of research including ours have shown that catechol groups in individual catechol-containing polyphenols and flavonoids play key roles in amyloid inhibition (Caruana et al, 2011; Sato et al, 2013; Velander et al, 2016; Wu et al, 2017). Extending from individual compounds, we recently demonstrated that catechols and redox-related quinones/anthraquinones are a broad class of amyloid inhibitors (Velander et al, 2020). The catechol moiety is a common structural component of many natural products, including scores of flavonoids and polyphenols, many of which were shown to have anti-diabetic, neuroprotective, and anti-aging activities (Cao & Raleigh, 2012; Ono et al, 2012; Yamada et al, 2015; Velander et al, 2017). RA, as a natural product containing dual catechol components, provides an outstanding example supporting our theory that catechol-containing compounds are amyloid inhibitors by way of demonstrating mechanistic additive effects. Catechol-containing flavonoids and polyphenols, with the advantages of low cost and few side effects as natural products, may have significant potential in translational applications as disease-preventating nutraceuticals.

RA is a natural compound identified from our library screening. With rational design using medicinal chemistry, more potent synthetic RA analog inhibitors may be engineered. The additive effect of RA provided an opportunity for us to further improve inhibitor potencies via a hybrid strategy. Our past work had demonstrated that such a strategy is effective in providing more potent analogs from initial natural product leads (Chojnacki et al, 2014; Liu et al, 2018). RA and its analog inhibitors will be useful not only for preclinical translational applications, but also as unique chemical biology probes in mechanistic studies, such as in understanding amylin amyloid-induced islet pathogenesis.

Orally administrated RA may present as glucuronide- or sulfate-conjugated or methylated forms in serum and it may be metabolized to conjugated forms of caffeic acid and ferulic acid (FA) (Baba et al, 2004). So active entities in serum after RA administration may include free and conjugated RA, methyl-RA, CA and FA. RA may have off-target effects or polypharmacology. For example, RA can interact with GABA transaminase and inhibit cyclooxygenase and acetylcholinesterase. While we did not observe any significant adverse effects in the HIP rat model, relevant biological responses may need to be evaluated when we design next-generation amylin amyloid inhibitors and proceed with expanded *in vivo* studies.

The mechanisms of amyloid inhibition by small molecules are not fully understood. This, and our past work led to several testable hypotheses: Does amylin amyloid inhibition by RA and other catechol-containing compounds occur via the mechanism of autoxidized *o*-quinone intermediates? Does RA conjugate with amylin through amine-groups of Lys^1^ or Arg^11^ or the amino terminal amine? We propose, as one potential mechanism, that the catechol functional groups in RA can be autoxidized to *o*-quinone intermediates and then covalently reacted with amylin to block amyloid growth (Velander et al, 2016; Velander et al, 2020). Quinones may be readily conjugated with the amine groups in Lys^1^ and/or Arg^11^ or at the amino terminal in human amylin and form Schiff base or Michael addition conjugates (Sato et al, 2013; Velander et al, 2016). We demonstrated such Schiff base conjugates in the case of another specific example of catechol-containing compound, baicalein (Velander et al, 2016). However, detection of such conjugates may be largely dependent on the conjugation efficiency of individual compounds. Such general covalent conjugation mechanisms, if validated, will have broad implications for both amyloid inhibitor design and mechanisms of inhibition for multiple amyloid systems in protein misfolding diseases. Our work, however, does not exclude other mechanisms, such as non-covalent inhibition mechanisms as proposed in the literature (Palhano et al, 2013; Tu et al, 2015; Velander et al, 2017).

Using EGCG as a benchmark, we provided strong evidence that both RA and control compound EGCG “remodeled” human amylin oligomers to form non-toxic amorphous aggregates, which had the characteristic broad range molecular weight “smears” in SDS-PAGE gel-based experiments (Figs. 5B & 5C). To further understand the mechanisms of inhibitor-induced amyloid remodeling, high-resolution characterizations of bound species, using approaches such as ion mobility spectrometry-mass spectrometry (Young et al, 2015), are necessary to define the interactions between RA and amylin.

Amylin amyloid deposition in the pancreas is well recognized a hallmark feature of T2D (Verchere et al, 1996; Hoppener et al, 2000; Westermark et al, 2011). Recently, amylin amyloid deposition was also found in the brains of T2D patients with dementia or Alzheimer’s disease, as well as in the brains of diabetic HIP animal models, raising the possibility that amylin amyloid may be a new amyloid in the brain (Jackson et al, 2013; Srodulski et al, 2014; Ly et al, 2017; Zhu et al, 2019; Despa et al, 2020). The exact mechanism of such diabetes-induced neurodegeneration remains elusive. However, validating this new hypothesis will be highly significant, as it opens a new field by identifying a specific molecular link between diabetes and neurodegeneration. Such validation will support the clinical and epidemiological observations that obese/diabetic individuals are more prone to dementia/neurodegeneration. RA, its analogues, or other effective amyloid inhibitors could provide useful chemical biology tools to validate this new hypothesis. It will be interesting to extend future investigations of amylin amyloid-induced pathology from the pancreas and diabetes to the brain and neurodegeneration.

## Materials and Methods

### Peptides, Chemicals, and Sera

Synthetic amidated human amylin was purchased from AnaSpec Inc. (Fremont, CA) and the peptide quality was further validated by the Virginia Tech Mass Spectrometry Incubator. Hexafluroisopropanol (HFIP) and thioflavin T (ThT) were purchased from Sigma Aldrich (St. Louis, MO). Rosmarinic acid, caffeic acid, and salvianic acid A were purchased from Toronto Research Chemicals (North York, ON, Canada), Fisher Scientific Inc. (Hampton, NH), and Sigma-Aldrich Corp. (St. Louis, MO) respectively. Dulbecco’s phosphate buffer saline (DBPS) pH 7.4, was purchased from Lonza (Walkersville, MD). Black 96-well non-stick-clear-bottom plates and optically clear sealing film were purchased from Greiner Bio-one (Frickenhausen, Germany) and Hampton Research (Aliso Viejo, CA) respectively. 300 mesh formvar-carbon-coated copper grids and uranyl acetate replacement solution were purchased from Electron Microscopy Sciences (Hatfeild, PA).

Lyophilized amylin powder (0.5 mg) was initially dissolved in 100% HFIP at a final concentration of 1-2 mM. The additional lyophilizing step was employed to eliminate traces of organic solvents, which have been shown to affect amylin aggregation. Aliquots were either lyophilized again prior to use in cell-based assays or dissolved directly into DPBS, 10 mM phosphate buffer pH 7.4 or 20 mM Tris-HCl pH 7.4 for all amylin amyloid-related *in vitro* assays. All remaining 1-2 mM stocks in 100% DMSO were stored at −80 °C until later use. The lyophilized powder from all compounds and ThT were dissolved in DMSO (10 mM) and distilled water or relevant buffer (1-4 mM). These stocks were stored at −20 °C until later use. Residual DMSO in the final samples used for all *in vitro* assays ranged from 0-9.5%. We determined that these DMSO concentrations had negligible effects on amylin amyloid aggregation as reflected by ThT fluorescence, TEM, Photo-Induced Crosslinking of Unmodified Proteins (PICUP) assays, and inhibitor-induced amyloid remodeling assays.

HIP rat sera and diabetic patient sera samples are generous gifts from Prof. Florian Despa (University of Kentucky). Detailed sample collection and institutional approval of usage were described in the literature (Despa et al, 2014; Srodulski et al, 2014).

### Cell Culture

Rat pancreatic INS-1 cells and mouse neuroblastoma Neuro2A cells were generously provided by Profs. Pierre Maechler (University of Geneva) and Deborah Good (Virginia Tech) respectively. The SH-SY5Y cells were purchased from ATCC (#CRL-2266). Vascular Smooth Muscle Cells (VSMCs) were primary cells isolated from mouse aorta using the treatment of type I, II collagenases and type III elastase. INS-1, Neuro2A, SHSY-5Y were cultured per vendor’s instructions. Briefly, INS-1 cells were cultured in RPMI-1640 medium containing 10% fetal bovine serum and supplemented with 11.1 mM glucose, 1 mM sodium pyruvate, 10 mM HEPES, 50 μM β-mercaptoethanol, 23.8 mM NaHCO_3_. The medium was changed every other day until the cells became confluent. Neuro2A cells were cultured in DMEM medium containing 1% non-essential amino acids, 1% L-glutamate and 10% FBS. SH-SY5Y cells were cultured in EMEM with 1% non-essential amino acids and 10% FBS. VSMCs were growing in DMEM containing 20% FBS.

### Cytotoxicity Assay and Inhibitor Activity Quantification

An MTT-based cell viability assay was used. The INS-1 cells, Neuro-2A, SH-SY5Y cells were seeded in 96-well plate at a density of 4×10^4^ cells/well. After 24 hours incubation, cells were treated with either human amylin (3.75 μM) with or without specified natural compounds. Following another 24 hours incubation, 0.9 mM MTT was added to each well. The reduced insoluble MTT formazan product was then dissolved in SDS-HCl lysis buffer (5 mM HCl, 5% SDS) at 37 °C. Cell viability was determined by measurement of absorbance change at 570 nm using a spectrometric plate reader. Peptides dissolved in buffer-treated cells were used as positive control and taken as 100%, and 0.5% Triton X-100 treated cells at the start of the incubation period with test peptides were used as negative control and taken as 0%.

### Thioflavin-T Fluorescence Assay

Fluorescence experiments were performed using a SpectraMax M5 plate reader (Molecular Devices, Sunnyvale, CA). All kinetic reads were taken at 25 °C in non-binding all black clear bottom Greiner 96-well plates covered with optically clear films and stirred for 10 seconds prior to each reading. ThT fluorescence was measured at 444 nm and 491 nm as excitation and emission wavelengths, respectively. Each kinetic assay consisted of final concentrations of 30 μM amylin and 10 μM ThT. The amount of time required to reach half maximum ThT intensity (t_1/2_) and inhibitory concentration (IC_50_) values for dose response curves were estimated by multi-parameter logistic nonlinear regression analysis. The transition from the lag-phase to the growth phase was estimated when the first measurable ThT fluorescence/time (slope) value exceeded ≥ 5-fold of the previous measured slope. All experiments were repeated three times using peptide stock solutions from the same lot. Natural product library used is as described (Velander et al, 2016). The NIH Clinical Collection (NIHCC) library contains a total of 700 FDA-approved drugs and investigational compounds with diverse chemical structures. The library was supplied in 96-well plates in DMSO. For screening, 384-well plate was used. Bravo liquid handler with 96-channel disposable tip head (Agilent Technologies, Wilmington, DE) was used to aid sample transfer. Total volume in each well was 15 μl. Residual DMSO (<5.4%) in each well had negligible effect on the fluorescent signals.

### Transmission Electron Microscopy (TEM) Analysis

TEM images were obtained by a JEOL 1400 microscope operating at 120 kV. Samples consisting of 30 μM amylin (20 mM Tris-HCl, 2% DMSO, pH 7.4) in the presence of drug or vehicle control were incubated for ≥48 hours at 37°C with agitation. Prior to imaging, 2-5 μL of sample were blotted on a 200 mesh formvar-carbon coated grid for 5 minutes and then stained with uranyl acetate (1%). Both sample and stain solutions were wicked dry (sample dried before addition of stain) by filter paper. Qualitative assessments of the amount of fibrils or oligomers observed were made by taking representative images following a careful survey of each grid.

### Gel-Based Remodeling Assay

Vehicle control or specified compounds were spiked into freshly dissolved amylin samples (containing amylin and buffer). Thereafter amylin aggregation was allowed to proceed for 3 days. Final amylin concentration was 15 μM that included 45 μM of compound (drug:amylin molar ratio was 3:1). After 3 days, these samples were vacuum dried and re-dissolved in 6.5 M urea containing 15 mM Tris and 1X SDS Laemmeli sample buffer, boiled at 95 °C for 5-10 minutes and subjected to SDS-PAGE followed by Western blot analysis with anti-amylin primary antibody (T-4157, 1:5000, Peninsula Laboratories, San Carlos, CA). All gel-based amyloid remodeling assays were repeated at least twice.

### Photo-Induced Crosslinking of Unmodified Proteins (PICUP) Assay

Amylin aliquots from a master mix in 10 mM phosphate buffer, pH 7.4 were added separately to 0.6 mL eppendorf tubes containing small molecule inhibitors or DMSO vehicle loaded controls. Crosslinking for each tube was subsequently initiated by adding tris(bipyridyl)Ru(II) complex (Ruby) and ammonium persulfate (APS) (Typical Amylin:Rubpy:APS ratios were fixed at 1:2:20, respectively, at a final volume of 15-20 μL), followed by exposure to visible light, emitted from a 150-Watt incandescent light bulb, from a distance of 5 cm and for a duration of 5 seconds. The reaction was quenched by addition of 1X SDS sample buffer. PICUP was visualized by SDS-PAGE (16% acrylamide gels containing 6 M urea), followed by silver staining. Final concentrations for all PICUP reactions included 30 μM of amylin and 150 μM of each compound.

### Animal Studies

HIP rats (RIP-HAT) and wild type Sprague-Dawley (SD) control rats were purchased from Charles River Laboratories (Wilmington, MA). Dietary supplementation approach was used to study the *in vivo* effects of RA on HIP rats amylin amyloid islet deposition, serum oligomer formation, and animal diabetic pathology. Dietary supplemental treatments were initiated when HIP rats were at prediabetic stage (5-6 month old) and the preventative treatment lasted for 4 months. Diet supplementation of RA (at a dosage of 0.5% w/w) was prepared by Envigo Inc. (Madison, WI). Bulk rosmarinic acid was purchased from Toronto Research Chemicals and its identity and purity were further validated at Virginia Tech Mass Spectrometry Incubator Facilities. Standard rodent chow diet (without RA incorporation) was used for the untreated HIP rats and wild type SD control rats groups. All animals were housed in temperature-controlled (23 ± 2°C) and light-controlled, pathogen-free animal quarters and were provided ad libitum access to diet. Animal protocol was approved by institutional IACUC committee. Sera samples for glucose and insulin analysis and amylin oligomer Western blotting analysis were collected in rat tail veins. Glucose levels were measured with an Agamatrix presto meter (AgaMatrix Inc., Salem, NH). Insulin levels were detected and quantified by a rat insulin ELISA kit (Mercodia AB, Uppsala, Sweden).

### Islet Immunohistochemistry and Amyloid Staining

At the end of animal studies, rats were euthanized and the pancreas tissues were dissected and fixed in 4% (v/v) formaldehyde buffer (pH 7.2). Pancreas samples were embedded in paraffin and sectioned by AML Laboratories Inc. (St. Augustine, FL). A series of tissue sections (5 μm thickness at 200 μm interval) were prepared, mounted on glass slides, and stained with Congo Red dye (Sigma Aldrich, St. Louis, MO) or with rabbit anti-amylin primary antibody (T-4157, Peninsula Laboratories, San Carlos, CA) followed by a rabbit ImmPRESS HRP anti-rabbit IgG (peroxidase) polymer detection kit and Vector NovaRED substrate kit (Vector Laboratories). Images were visualized by microscopy (Zeiss Axio Observer, Thornwood, NY) or Nikon Eclipase Ti-U and were analyzed with Zeiss 2 (blue edition) or NIS-Elements AR 411.00. Programs.

### Statistical Analysis

All data are presented as the mean ± S.E.M and the differences were analyzed with a one-way analysis of variance followed by Holm-Sidak’s multiple comparisons (amylin kinetics) or unpaired Student’s *t* test. These tests were implemented within GraphPad Prism software (version 6.0). *p* values < 0.05 were considered significant.

### MD Simulations

*Starting Structure* In order to simulate trimer formation of amylin in a reasonable time scale using MD simulations, the principal amyloidogenic region (residues 20-29) was extracted from an equilibrated MD structure of full length (residues 1-37) human amylin (PDB access code 2L86; Nanga et al, 2011). This PDB structure was chosen to best represent biologically active, monomeric amylin given the presence of the disulfide bound between Cys^2^-Cys^7^ and C-terminal amidation. However, 2L86 was solved in an SDS micelle environment and to better mimic experimental procedures, we sought to use a structure of amylin that mimicked solution conditions (water and 150 mM NaCl). Simulations of full-length amylin were constructed by placing the structure of 2L86 in a cubic box with a minimum solute-box distance of 1.0 nm. The SPC water model (Berendsen et al, 1981) was used to solvate the box, and Na^+^ and Cl^-^ ions were added to reach a final concentration of 150 mM NaCl and maintain a net neutral system. All simulations were done using the GROMACS software package, version 4.6.0 (Hess et al, 2008; Pronk et al, 2013) and the GROMOS53a6 force field (Oostenbrink et al, 2004). Energy minimization was performed using the steepest descent method, with all protein heavy atoms being restrained during equilibration and later released during MD simulation. Equilibration was performed in two sequential steps, NVT and NPT. Three replicates starting with different random starting velocities in NVT were performed. NVT was performed for 100 ps and maintained at 310 K using the Berendsen weak coupling method (Berendsen et al, 1984). Following NVT, NPT dynamics using a Nosé-Hoover thermostat (Nose, 2002; Hoover, 1985) and a Parrinello-Rahman barostat (Parrinello & Rahman, 1981; Nose, 1983) to maintain temperature (310 K), and pressure (1 bar) was performed for 100 ps. Following equilibration, MD simulations were run using three-dimensional periodic boundary conditions, with short-range cutoffs of 1.4 nm being applied to all non-bonded interactions. Long-range interactions were calculated using the smooth particle mesh Ewald (PME method; Darden et al, 1993; Essmann et al, 1995) using cubic interpolation and Fourier grid spacing of 0.16 nm. Bond lengths were constrained by P-LINCS (Hess, 2008) using an integration time step of 0.2 fs. Simulations were run until the criteria of convergence was met, which consisted of stabilization of 100 ns for both backbone root-mean-square deviation (RMSD) and protein secondary structure, as determined by the DSSP algorithm (Kabsch & Sander, 1983). RMSD clustering was used to produce a representative structure of the last 100 ns of each replicate (Daura et al, 1999). The center structure of the replicate cluster structure whose average secondary structure was closest to the set average was chosen as the template for simulations of trimer formation of amylin_(20-29)_.

*Trimer Formation* Residues 20-29 of amylin were extracted from the representative structure generated by MD simulations of full-length amylin in water and 150 mM NaCl. This was done in order to have native, monomeric fragments of this region rather than the preformed fibrils. Acetyl and amide groups were added to cap the ends of the ten-residue fragment to negate spurious effects from having charges at the termini. The sequence simulated of the amylin_(20-29)_ fragment is as follows: Ace-SNNFGAILSS-NH2. Three amylin_(20-29)_ fragments were randomly placed at least 2.0 nm away from each other in a cubic box with dimensions of 11 x 11 x 11 nm, with a solute-box distance of 1.0 nm. For control simulations, no small molecules were added to the system. For simulations with rosmarinic acid, caffeic acid, and salvianic acid A, five molecules were randomly placed in the system at least 2.0 nm away from amylin_(20-29)_ fragments and other small molecules. Topologies for rosmarinic acid, caffeic acid, and salvianic acid A were generated using the PRODRG2 server (Schuttelkopf & van Aalten, 2004) and refined by using charges and atom types found in functional groups within the GROMOS53a6 force field. For the ester functional group, parameters were taken from literature (Horta et al, 2011) and modified to account for the double bond in the “R” position. Parameters and charges for all small molecules used in this study can be found in the supplemental information. RA, CA, and SAA were deprotonated to mimic their protonation state at pH 7.4 (net charge −1). The following systems were generated and are identified in the following way: Control – 3 amylin_(20-29)_ fragments and no small molecules, RA – 3 amylin_(20-29)_ fragments and five molecules of rosmarinic acid, CA -- 3 amylin_(20-29)_ fragments and five molecules of caffeic acid, and SAA – 3 amylin_(20-29)_ fragments and five molecules of salvianic acid A. These systems were then equilibrated and were run in MD simulations in the same way as full-length amylin. Simulations were run to 600 ns, which was found to be an adequate amount of time for trimer formation and ß-strand structure to remain stabilized in the control simulations. Backbone RMSD and secondary structure analysis were also used to determine convergence. Analysis was performed using programs in the GROMACS package and over the last 100 ns of simulation time (e.g., 500-600 ns) in order to report on stabilized trimer structures or what their corresponding trimer structure looks like in the presence of small molecules. RMSD clustering with a cutoff of 0.2 nm was used to generate the structures. PyMOL was used for molecular visualization and statistical analysis was done using a two-tailed *t*-test, with significance determined if *p* < 0.05.

## Acknowledgments

This work was supported in part by the Hatch Program of the National Institute of Food and Agriculture, USDA (BX and DRB), Commonwealth Health Research Board Grant 208-01-16 (BX, DRB, and SZ), NIH grants R03AG061531 (BX) and R01AG058673 (SZ), Diabetes Action Research and Education Foundation Grant (BX), Awards No. 16-1, 18-2 and 18-4 from the Commonwealth of Virginia’s Alzheimer’s and Related Diseases Research Award Fund (BX, LW, and SZ), Alzheimer’s Association/Michael J. Fox Foundation grant BAND-19-614848 (BX), and Alzheimer’s Drug Discovery Foundation 20150601 (SZ). We thank Prof. Florian Despa (University of Kentucky) for generously providing HIP rat and diabetic patient sera samples. We thank Ms. Kathy Lowe at Virginia-Maryland Regional College of Veterinary Medicine for her excellent technical assistance in collecting TEM data.

## Author Contributions

L.W. and P.V. performed majority of the experiments, acquired and analyzed the data. A.M.B. assisted with the molecular simulation studies and analyzed the data. Y.W. assisted with the diabetic animal studies and tissue analyses. D.L. contributed to the design of animal studies and diabetic animal characterization. D.R.B. designed the molecular simulation experiments and analyzed the data. S.Z. contributed to drug discovery experimental design. B.X. conceived, organized, designed the experiments, and analyzed the data. B.X., L.W., P.V., A.B., D.R.B. wrote the paper.

## Additional Files

Supplementary Files

Three supplementary tables and two supplementary figures are attached.

## Supplemental Data

**Table S1.** Average secondary structure (shown in %) of trimer fragment formation (residues 20 - 29) of amylin with or without the presence of small molecules.^*a*^

| System | Coil | β-strand | α-helix |
| --- | --- | --- | --- |
| WT | 42 ± 11 | 58 ± 11 | 0 ± 0 |
| RA | 73 ± 13 | 28 ± 13 | 0 ± 0 |
| CA | 63 ± 17 | 37 ± 17 | 0 ± 0 |
| SAA | 52 ± 17 | 45 ± 20 | 2 ± 3 |

<sup>a</sup> Percentages represent averages of 3 replicates over the final 100 ns of simulation time, with associated standard deviations.

**Table S2.** Average hydrogen bond presence between amylin fragment peptides (20-29), amylin fragment peptides and SMs, and between SMs.^*a,b*^

| System | Peptide-Peptide | Peptide-SM | SM-SM |
| --- | --- | --- | --- |
| WT | 19 ± 3 | n/a | n/a |
| RA | 12 ± 3 | 13 ± 4 | 3 ± 1 |
| CA | 15 ± 2 | 4 ± 4 | 0.1 ± 0.1 |
| SAA | 17 ± 1 | 0.7 ± 0.3 | 0.1 ± 0.1 |

<sup>a</sup> Averages of 3 replicates over the final 100 ns of simulation time, with associated standard deviations. SM: Small Molecule.
<sup>b</sup> n/a represents not applicable to the system tested.

**Table S3.** Average total SASA (nm^2^) of amylin fragments (20-29) and SMs.^*a,b*^

| System | Peptide | SM |
| --- | --- | --- |
| WT | 24 ± 1 | n/a |
| RA | 29 ± 2 | 19 ± 1.6 |
| CA | 26 ± 1 | 14.7 ± 0.1 |
| SAA | 26 ± 1 | 13.9 ± 0.1 |

<sup>a</sup> Averages of 3 replicates over the final 100 ns of simulation time, with associated standard deviations.
<sup>b</sup> SASA: solvent accessible surface area. n/a represents not applicable to the system tested.

**Figure S1.**
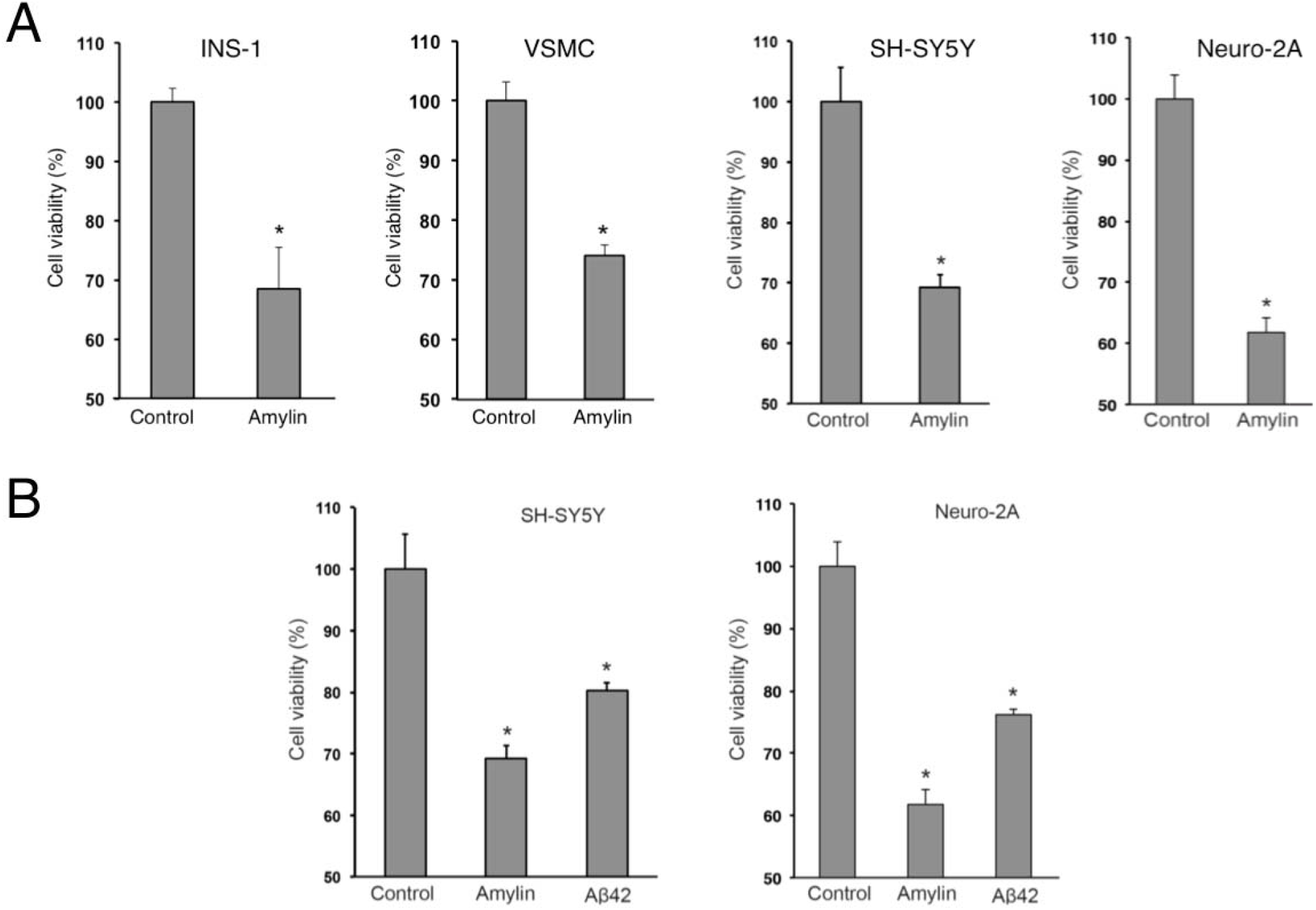
Cytotoxicity of amylin amyloid on various cell lines and comparison of cytotoxicity of amylin amyloid versus Aβ amyloids. (A) Cytotoxicity assays demonstrate the toxic effects of amylin against multiple cell lines relevant to the pancreas, heart, and brain: pancreatic β-cells INS-1, mouse vascular smooth muscle cells (VSMCs), human SH-SY5Y neuronal cells, and mouse Neuro-2A cells, each at 5 μM concentration. (B) Neurotoxicity assays demonstrate that amylin amyloid is more toxic than a well-established control, amyloid β-peptide Aβ42. Concentrations for each peptide were 10 μM for SH-SY5Y treatment or 15 μM for Neuro2A treatment.

**Figure S2.**
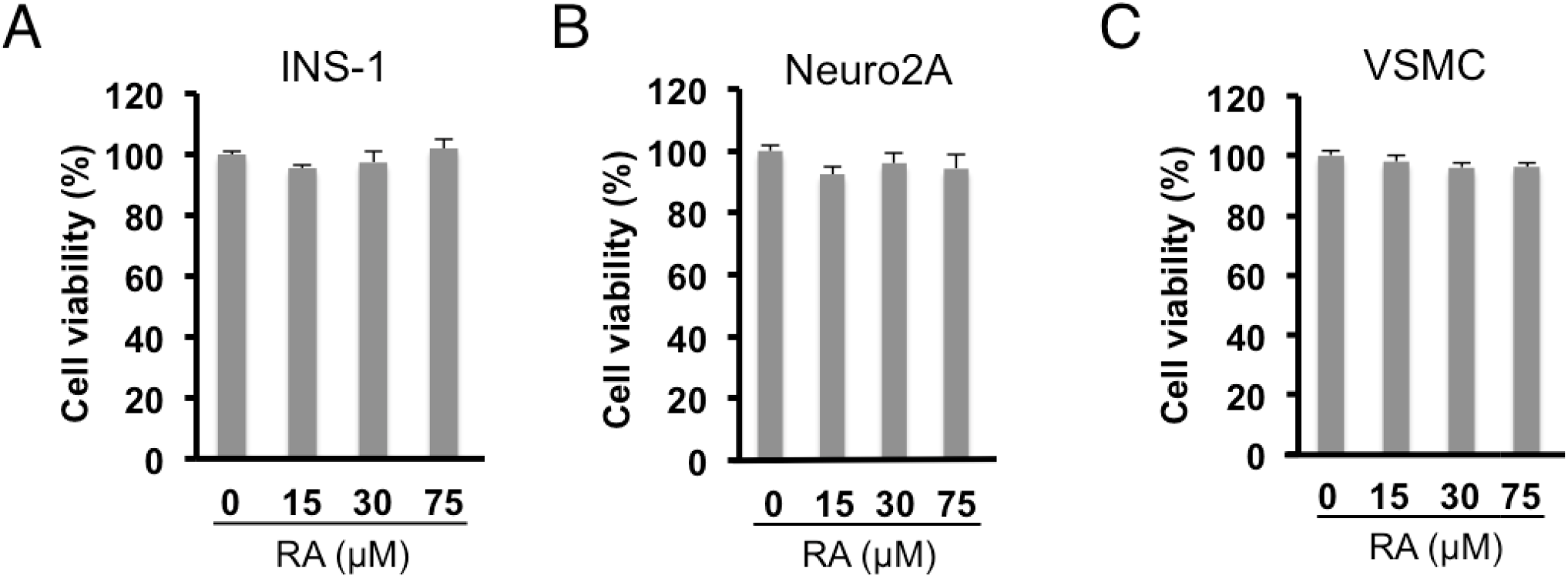
Control experiments demonstrate that RA has no effects on cell viability at the specified concentrations in INS-1 (A), Neuro2A (B), and VSMC cell lines (C). Concentration of amylin was 15 μM in each case.

